# Diffusion Probabilistic Models for Missing-Wedge Correction in Cryo-Electron Tomography

**DOI:** 10.64898/2026.02.15.706025

**Authors:** Nadeer Hasan, Aurélie Bertin, Slavica Jonic

**Affiliations:** IMPMC-UMR 7590 CNRS, Sorbonne Université, MNHN, Paris, France; PCC-CNRS UMR 168, Institut Curie, Sorbonne Université, Paris Sciences et Lettres, Paris, France

**Keywords:** Cryo-electron tomography (cryo-ET), Missing Wedge, Diffusion Probabilistic models

## Abstract

Interpretation of 3D cryo-electron tomography (cryo-ET) reconstructions (tomograms) is hampered by the so-called missing-wedge (MW) distortions, which arise because tilt image series used for the reconstructions are acquired in a limited angular range. While many deep-learning approaches address the correction of the MW artifacts on the level of tomograms (3D volumes), the correction at the level of 2D tilt images (generation of unacquired images) remains underexplored. We propose MW-RaMViD, a 2D tilt-image generation method for MW correction, based on Random-Mask Video Diffusion (RaMViD) method for prediction of frames in natural videos. To adapt RaMViD for cryo-ET, we add MRC image-format support, floating-point pixel intensity representation, and a controlled inference protocol enabling both one-run and progressive MW completion (generating a small number of missing tilts per step using a sliding window). We evaluate the method on a synthetic noisy tilt-series dataset and study the effects of MW completion step size and conditioning sequence length. Results show that smaller step sizes and larger conditioning windows reduce error accumulation at higher tilt angles and improve reconstruction fidelity, which was measured by Root Mean Square Error on the image level and by Fourier Shell Correlation on the tomogram level.

## I. Introduction

Cryogenic electron tomography (cryo-ET) has rapidly advanced as a central method in structural biology for visualizing biological specimens in near-native states, enabling three-dimensional (3D) imaging of cellular architecture and the spatial organization of proteins and organelles at nanometer-scale resolution, without requiring crystallization or staining. Cryo-ET relies on tilting the sample incrementally in a limited angular range (usually [−60°, 60°] and sometimes [−45°, 45°]), with an angular step (usually 1°, 2°, or 3°), and collecting a two-dimensional (2D) projection image at each tilt angle [1]. From the collected tilt-image series (the so-called tilt series), a 3D volume (the so-called tomogram) can then be calculated using classical or deep learning-based methods [2] [3].

During tilting in the electron microscope, the same sample is exposed to electrons multiple times and accumulates radiation damage. To preserve the structure of biological samples, they are exposed to low electron doses, which induces a low signal-to-noise ratio (SNR) of the collected data [4]. Additionally, as the effective sample thickness increases at higher tilt angles, the electrons travel longer paths through the sample, which increases electron diffusion and induces image blurring. At very high tilt angles, the increased sample damage and image blurring, in addition to a low SNR, would make the data collection useless [5]. Therefore, such high tilt-angle data are not acquired. As a result, a substantial portion of angular information is missing and produces an empty, wedge-shaped region on the 3D reconstructed tomogram in Fourier space. This region is known as the missing wedge (MW) and induces artefacts on the tomogram in real space (notably, elongations in the direction perpendicular to the tilt axis) [6] [7]. Such distortions hamper deciphering of fine structural details of biological samples (Figure 1).

**Figure 1.**
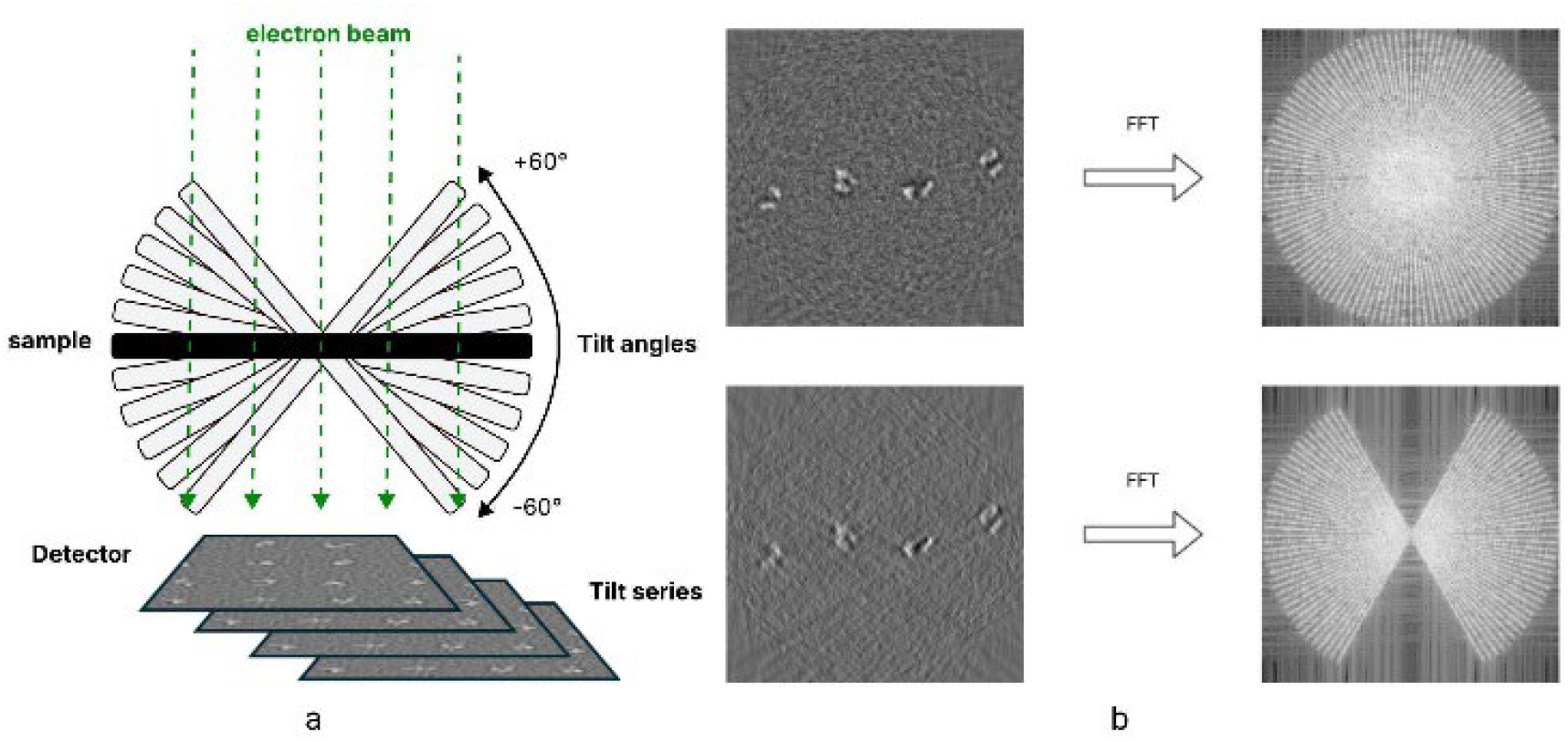
Cryo-ET data collection geometry and missing wedge (MW). (a) Tilt-series data collection. (b) Central slice of a tomogram (left) and 2D Fourier transform of this slice (right) for 3D reconstructions from images synthesized by tilting a synthetic sample.

Recently, deep learning methods have been proposed for MW correction. Most of them address the MW problem using supervised or self-supervised learning on volumetric data, typically on volumes extracted from tomograms (subtomograms), such as IsoNet [7] and DeepDeWedge [8]. Other approaches start from the collected tilt series but still learn a 3D representation (e.g., via latent back-projection) [9]. In contrast, methods that predict missing information in 2D real space, directly from the tilt series, remain underexplored. Recently, cryoTIGER [10] showed that methods designed for prediction of frames in natural videos can be used for cryo-ET tilt-image prediction. However, CryoTIGER improves sampling in the angular range used for data collection, by interpolating intermediate tilt images with deep frame-interpolation network (FILM) [11], but it does not allow to generate tilt images in the MW region.

In this article, we propose an alternative 2D approach that is based on probabilistic diffusion model called Random-Mask Video Diffusion (RaMViD), originally developed for video prediction and infilling [12]. We consider a tilt series as a sequence of past video frames to predict future frames, namely tilt images in the MW region. This is possible as adjacent tilt images are not independent but consecutive projections of the same sample observed under gradually changing viewing geometry [1]. Our method (named MW-RaMViD) performs MW correction prior to 3D reconstruction, by tilt-image synthesis in the MW region. Because ground-truth images in the MW region are unavailable in practice, MW-RaMViD was validated in a controlled setting using synthetic noisy tilt series. Our experiments show that MW-RaMViD produces tomograms of improved quality, with strongly reduced MW artifacts. To the best of our knowledge, MW-RaMViD is the first diffusion-based approach for MW correction.

The article is further organized in five sections. Section II recalls the basic principles of diffusion models and RaMViD. Section III presents our work performed to adapt RaMViD to cryo-ET MW correction. In section IV, we describe data synthesis and experiments performed with this data using MW-RaMViD, and discuss results. We conclude the article in section V.

## II. Background

In generative modelling, neural networks are trained to capture underlying data distributions. Two common families are implicit generative models [13], such as Generative Adversarial Networks GANs [14], which generate samples without an explicit likelihood, and likelihood-based models, such as Variational Autoencoders VAEs [15], which define and optimize a probabilistic model. In practice, GANs can suffer from training instability or mode collapse [16], while likelihood-based models often rely on simplifying assumptions that can limit sample fidelity [17]. Score-based generative modelling offers an alternative, where instead of modelling data density directly, it learns a score function (gradient of log-density ∇_*x*_ log *p* (*x*)) across multiple noise levels, enabling sampling by iteratively denoising from a simple prior [18]. Among score-based methods, diffusion probabilistic models have emerged as a particularly effective approach, combining a tractable forward corruption process with a learned reverse-time generative model.

### A. Diffusion process

Diffusion models construct data through a two-step process: a forward diffusion that gradually corrupts input data by adding Gaussian noise over many time steps, transforming the data distribution into a simple prior, and a learned reverse process implemented by a spatiotemporal neural network (typically a U-Net denoiser) that denoises and reconstructs samples from noise [19]. In the score-based formulation, a clean sample *x*_0_ ∈ *R*^*d*^ is drawn from the data distribution *p*_data_(*x*_0_) and perturbed over diffusion steps *t* ∈ {0, …, *T*} using a noise schedule σ(*t*) producing noisy variables *x*_*t*_ [18]. The forward corruption has the closed form:

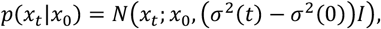

which can be sampled using the reparameterization trick

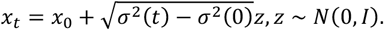

This parameterization makes it possible to define a time-conditional score network *s*_*θ*_(*x*_*t*_, *t*) trained by denoising score matching (DSM) to estimate the score across noise levels:

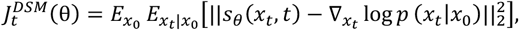

where the target score is available in the closed form [18] [20]:

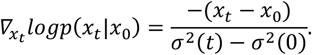

In many applications, including our tilt-series completion setting, the goal is not sampling from *p*(*x*) but from a conditional distribution *p*(*x*|*y*) where *y* is an additional information such as class, text or (most relevant here) a known subset of the data to be completed. Two common conditioning strategies are the Conditional Diffuse Estimator (CDiffE) [21], which trains an unconditional score model and introduces conditioning only during sampling, and Conditional Denoising Estimators (CDE) [21] [22], which train a conditional score model *s*_*θ*_(*x*_*t*_, *y, t*) directly..

### B. Random-Mask Video Diffusion (RaMViD)

RaMViD adapts score-based diffusion to conditional sequence generation by introducing conditioning through masking. Given a sequence *x*_0_ ∈ *R*^*L*×*H*×*W*×*P*^ of length *L*, where *H*and *W* are the height and width of each frame, and *P* is the number of channels, RaMViD partitions the frame indices into a conditioning set *C* and an unknown set *U*, where *U* ∩ *C* = Ø, *U* ∪ *C* = {0, …, *L* − 1}, and 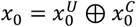. For diffusion steps *t* ∈ {0, …, *T*}, only the unknown frames are corrupted, while the conditioning frames remain clean:

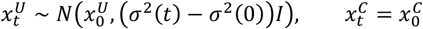

So, the input to the denoising network (U-Net) is the full mixed sequence 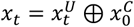 that contains noisy frames at indices in *U* and clean frames at indices in *C*. The score network *s*_*θ*_(*x*_*t*_, *t*) is then trained with a denoising score-matching objective computed only on the unknown part *U* denoted as 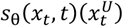 [12],

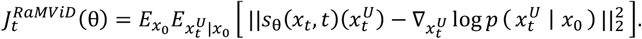

This design forces the model to learn how to reconstruct the missing frames while using the conditioning frames as a context at every diffusion step, which improves consistency between generated and observed parts of the sequence.

A second key feature of RaMViD is randomized masking. During training, the number and indices of conditioning frames in the training sequence are sampled randomly (up to a value of *K*, which is a hyperparameter). So, the same model learns to handle many completion and infilling patterns. At inference time, the conditioning set *C* is fixed to the available frames and the model generates the remaining frames in *U* by running the reverse diffusion process [12].

## III. MW-RaMViD: CRYO-ET EXTENSION OF RaMViD

In our formulation of MW correction, a tilt series is a video frame sequence. Our goal is to synthesize uncollected tilt images and then perform a tomographic reconstruction. To achieve this, we adapt RaMViD (Section II) for MW correction. RaMViD was originally designed and evaluated on natural video datasets, where inputs are RGB clips stored in standard formats (e.g., MP4/AVI). To apply this framework to cryo-ET tilt series, several adaptations were required, including enabling the readout of tilt-series data, commonly available as stacks of images in MRC floating-point format, and a full control of sampling to enable a precise order of tilt generation. In particular, we adapted the original data reading and preprocessing (input 8-bit integer data) to floating-point MRC data format, which preserves the cryo-ET data intensity and noise statistics relevant for downstream analyses. Regarding data preprocessing, RaMViD rescales 8-bit video intensities to [−1,1], while for floating-point MRC tilt series, we implemented per-series min–max normalization to [−1,1]. Besides, because MW completion targets specific missing tilt angles, we implemented controlled evaluation and progressive completion protocols. In particular, we enabled a consistent and progressive tilt-angle extrapolation via sliding-window generation with a chosen stride to generate a specified number of missing tilts per step.

## IV. Experiments

### A. Data

To evaluate the performance of MW-RaMViD under controlled conditions, we synthesized tilt-series data from a simulated sample containing multiple instances of a protein in motion (different frozen conformations), with added noise and contrast transfer function (CTF) of the microscope. Chain A of adenylate kinase from the Protein Data Bank (PDB) structure 4AKE (referred to as AK) was used as the initial structure for simulating different conformations. The motion was simulated using a linear combination of normal modes (vectors describing frequency-depending harmonic motions). As in similar studies [23] [24] with simulated protein dynamics data, motion was synthesized using lowest-frequency non-rigid modes as they describe large-scale, biologically relevant motions. Using ContinuousFlex software [25], we generated diverse conformations of AK by sampling random linear combinations of modes 7 and 8, with uniform normal-mode amplitude distribution (it should be noted that the first six modes describe rigid-body motions). Then, 300 volumes of size 256×256×100 voxels were generated (voxel size: 2.2 A) to emulate a thin section of a cell (thickness: 100 voxels, that is 22 nm), each with 20 instances of AK in random conformations, orientations, and positions (random shifts with respect to a regular grid used to partition the volume in 20 boxes, each containing one instance of AK).

Two types of tilt-series datasets were then obtained using these 300 volumes, one for the training and the other for the evaluation of sampling. To this goal, each volume was tilted from −90° to +90° with an increment of 3° and, for each tilt, projected on a static image plane. This resulted in 300 simulated tilt series, each with 61 tilt images of size 256×256 pixels. Then, the images were degraded with CTF and noise (0.5 μm defocus, SNR ≈ 0.1) to simulate experimental conditions. The complete tilt-series (61 tilt images) were used only for evaluation of the inference (sampling), whereas the training was performed using only tilt images from the range [−60°, +60°] (only 41 tilt images). The projections outside the range [−60°, +60°] (20 tilt images) were not used for training to mimic the practical setting where such high-tilt images are not collected, but they were used to evaluate the results of the MW inference (as the ground truth). The training was performed using only 200 out of 300 simulated tilt-series. The remaining tilt series were used for inference.

### B. Training procedure

We trained MW-RaMViD using multiple GPUs in parallel on the synthetic data described above. The diffusion process corrupts images using a cosine noise schedule with *T*=50,000 diffusion steps (50,000 discrete noise levels). The reverse (denoising) process uses a multi-resolution U-Net with one residual block per resolution level and 128 channels at the input of the first block. Optimization is performed with AdamW at a learning rate of 2×10^-5^. An exponential moving average (EMA) of the model parameters is maintained during training (EMA decay 0.9999), and we use the EMA checkpoint for sampling and evaluation.

Training was run for 70,000 optimizer updates using 4 GPUs, with a batch size of 1 per GPU, which corresponds to a sequence of length *L*=8 extracted from a tilt series. Therefore, at each update, 4 sequences of length *L*=8 are processed in parallel and gradients are synchronized to perform one global update. Furthermore, at each update, for each sequence, a level of noise is sampled from the entire set of *T*=50,000 noise levels (rather than simulating the full *T*-step forward diffusion trajectory) and this noise is applied to the unknown part of the sequence. Within the sequence, up to 4 frames are treated as unknown (the frames that will be corrupted with noise), while the remaining frames are kept clean as conditioning context (conditioning frames that will not be corrupted).

In the remaining part of the article, the known (conditioning) frames will be referred to as given frames and their number will be represented by *g*, whereas the unknown frames will be referred to as predicted frames and their number at one run will be represented by s, with *L=g+s*. The length *L* can have different values for training and sampling (inference).

### C. Sampling strategies

We designed several MW completion strategies to study the effects of g and s. In all cases, the given frames immediately precede the predicted frames. The following three types of strategies were compared: 1) *s=*20, i.e., a simultaneous completion of all 20 missing tilt angles (10 positive and 10 negative); 2) *s=*10, i.e., a simultaneous completion of 10 positive or 10 negative missing angles; 3) a progressive, step-by-step completion with a step *s* and a sliding window of length *g* that moves over the tilt series in each step to include *s* newly generated frames into the given frames for the next step.

### D. Evaluation metrics

In this article, we perform qualitative and quantitative evaluations of the results of the MW completion using the different sampling strategies described in subsection *C*. For the quantitative evaluation, we use two metrics. One is the root mean squared error (RMSE) between the predicted and ground-truth tilt images, with images normalized using min–max normalization before the RMSE calculation (a min–max normalization was used with the min and max values obtained from the ground-truth intensity range of the predicted frames). The second evaluation metric is the Fourier Shell Correlation (FSC) between two tomograms: one reconstructed from a tilt series consisting of a mixture of known images (in the range [−60°, 60°]) and predicted images (outside the range [−60°, 60°]) (here, referred to as predicted tomogram), and the other (here, referred to as ground-truth tomogram) reconstructed from the complete ground-truth tilt series (known images in the range [−90°, 90°]).

The FSC between two volumes is obtained as a function of spatial frequency:

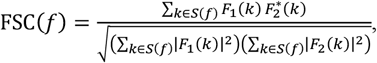

where *F*_1_ and *F*_2_ are the Fourier transforms of two volumes and *S*(*f*) denotes a Fourier shell at spatial frequency *f*.

Ideally, the FSC curve should be 1 at all frequencies, which would mean two identical volumes. As a baseline for comparing the FSC curves for different sampling strategies, we use the FSC between the ground-truth tomogram and a tomogram reconstructed using only known images (range [−60°, 60°]) and no MW correction (here, referred to as MW-affected tomogram). This baseline FSC is used to show the difference between the FSC before and after MW correction using different MW completion strategies. We expect an increase in the FSC at all frequencies after the MW correction. All tomograms were reconstructed in Scipion [26] using fast Fourier reconstruction based on central slice theorem.

### E. Results

The RMSE and FSC results shown in this section and all figures (Figures 2-5) were obtained for a single tilt series from the set of 100 tilt series used for the inference. Similar results were observed for 10 other tilt series randomly selected from the inference dataset.

**Figure 2.**
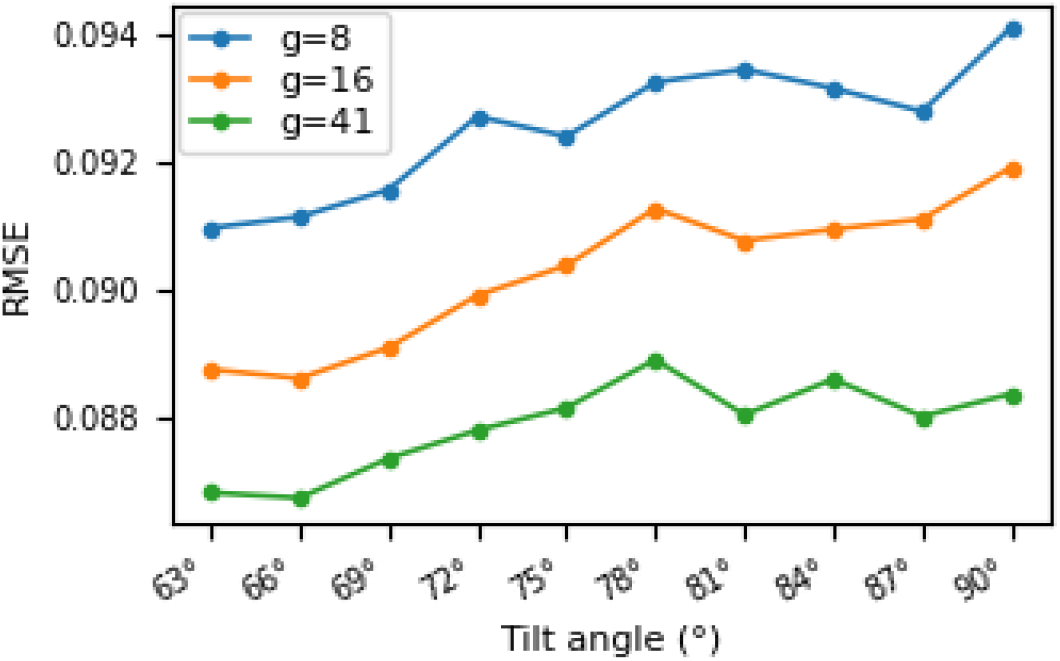
RMSE between the predicted and ground-truth frames for positive tilt angles (from +63° to +90°) when varying the number of given frames g. Similar results were obtained for negative tilt angles (unshown data).

To study the effect of conditioning length *g*, we varied *g* while keeping *s* constant. The RMSE results for *s=*2 and *g*∈{8,16,41} show the best prediction for *g=*41 (Figure 2). Therefore, we performed the tests of the three MW completion strategies described in subsection *C* (*s=*20, *s=*10, and step-by-step) using *g=*41. For the step-by-step completion strategy, we performed the experiments using the following values for step *s*: *s*∈{5,4,3,2,1}. Across different experiments, we obtained similar results for positive and negative tilt angles, but for sake of space, we only show the results for positive angles.

Figure 3 shows qualitatively a gradual degradation of the prediction quality for *s=*20 starting at about the fifth predicted frame (75°) and a better preservation of the structural content in these frames for *s=*1. The RMSE further confirms that smaller steps *s* lead to globally better prediction quality (Figure 4), showing that the best results are obtained with *s=1*.

**Figure 3.**
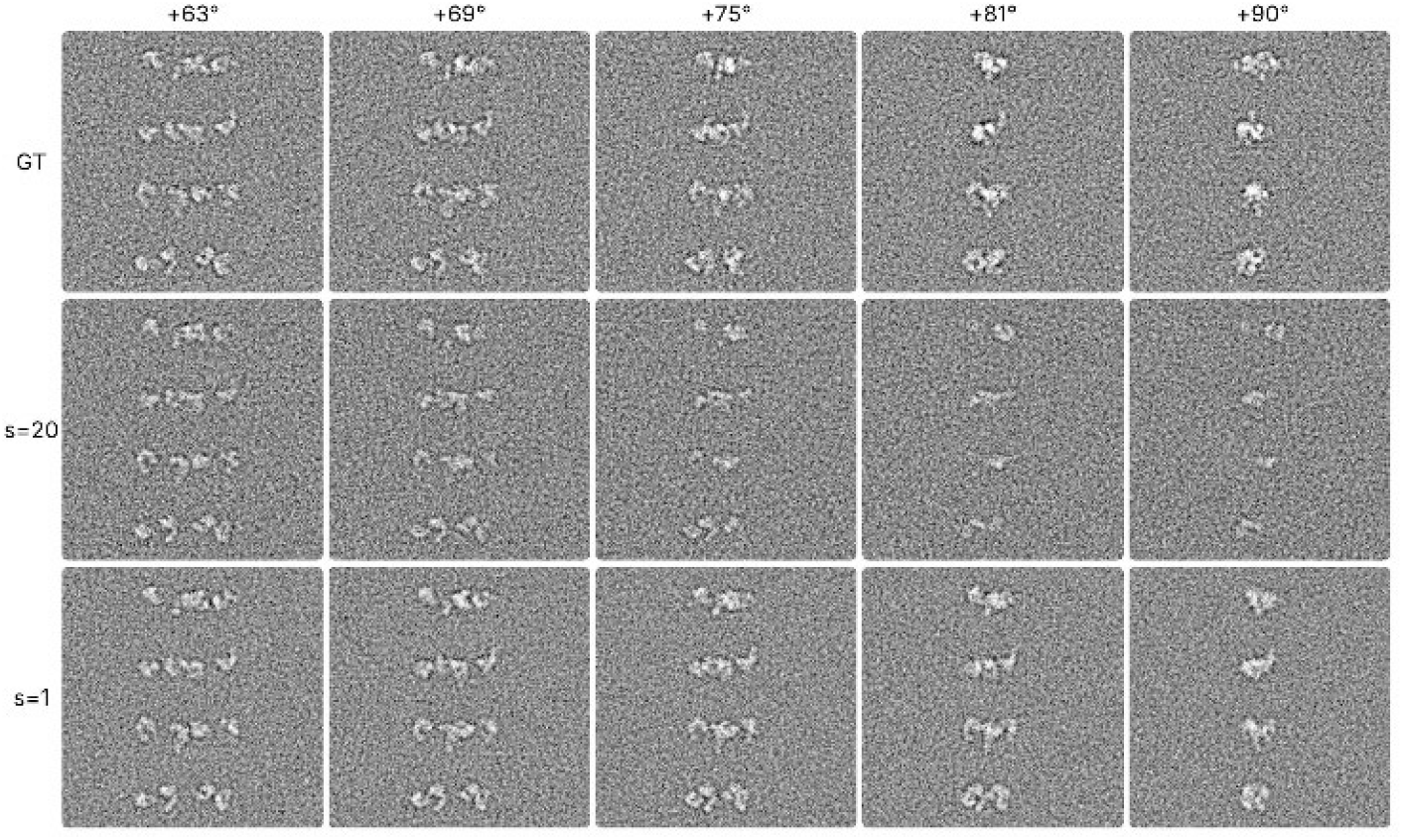
Qualitative comparison of predicted and ground-truth frames for positive tilt angles and two step-by-step MW completion strategies (s=20 and s=1). Columns from left to right: 63°, 69°, 75°, 81°, 90°. First row: ground truth. Second row: s=20. Third row: s=1.

**Figure 4.**
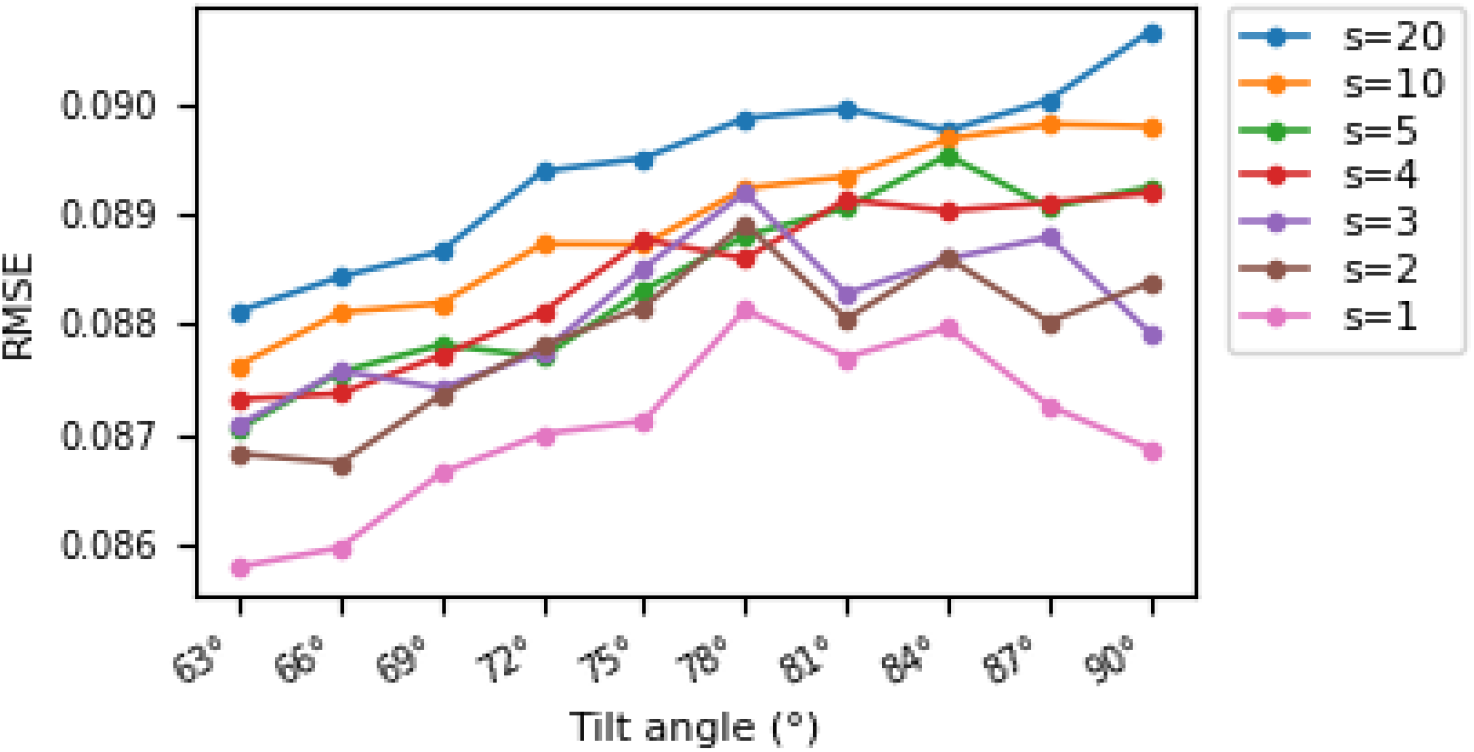
RMSE between the predicted and ground truth frames for different step sizes s and positive tilt angles (from +63° to +90°). Similar results were obtained for negative tilt angles (unshown data).

In Figure 5, the baseline FSC shows low correlation between the MW-affected and ground-truth tomograms at high frequencies. It should be noted that this low correlation is due to the missing wedge, because the MW-affected tomogram was reconstructed without the high-tilt images that are present in the ground-truth tomogram (images in the angular ranges [−90°,−60°] and [60°,90°]). On the other hand, the predicted tomograms include synthesized images (using our sampling strategies) in these missing angular range. For all values of *s* and across the entire frequency range, the FSC between the predicted and ground-truth tomograms has higher values than the baseline FSC, which indicates the MW correction. The differences between sampling strategies are most evident at low spatial frequencies (inset in Figure 5), where *s=*1 achieves the highest FSC values indicating the best MW correction results.

**Figure 5.**
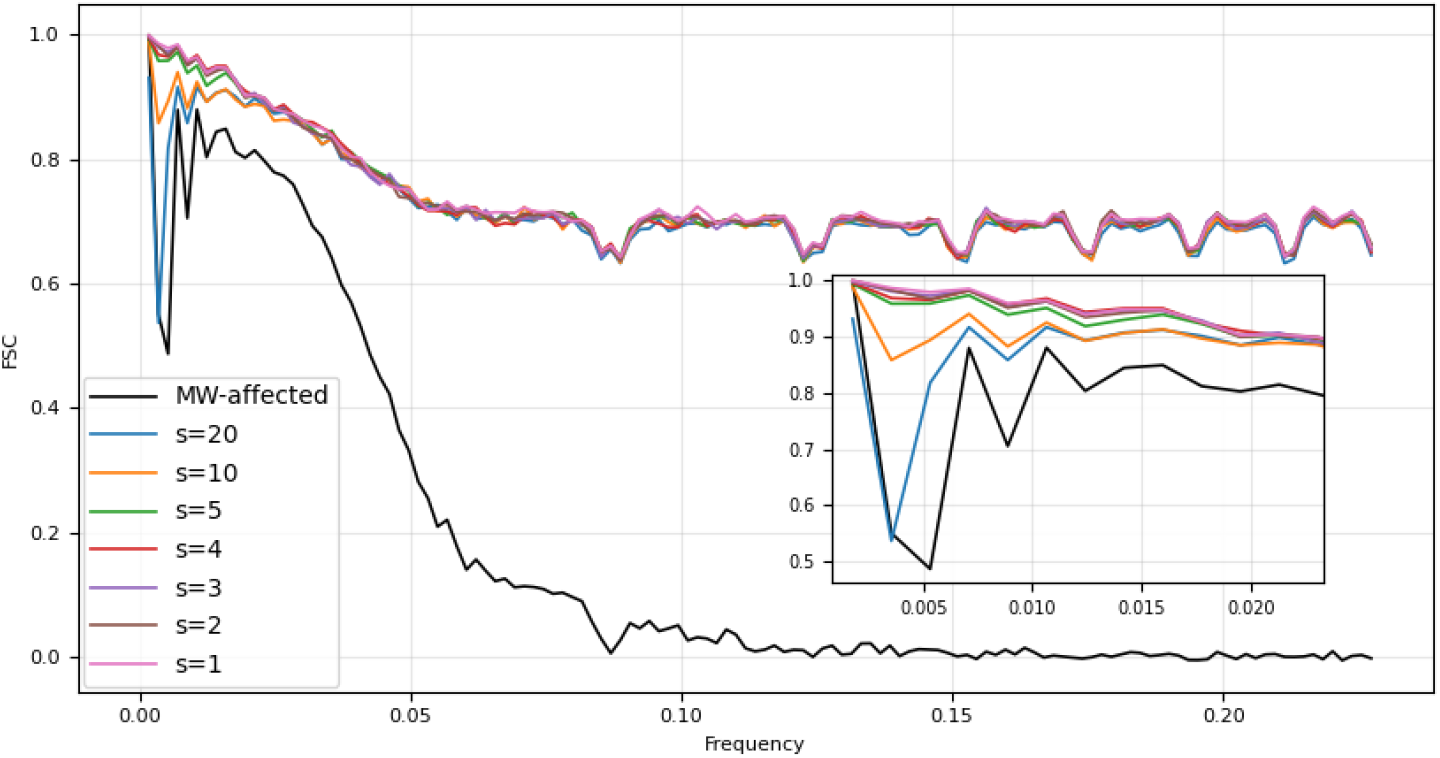
Superposition of the FSC curves calculated between the ground-truth and predicted (for different s) tomograms and the baseline FSC curve calculated between the MW-affected and ground-truth tomograms (black line indicated in the legend as “MW-affected”). Inset: zoom into the low-frequency region.

### F. Speed

Training was performed on 4 NVIDIA V100 GPUs (32 GB VRAM), which required about 50 wall-clock hours for 70,000 optimization steps. Inference (sampling) was performed on a single NVIDIA V100 GPU (where 10,000 steps of reverse diffusion process were executed). The inference time increases with the sequence length *L*. For instance, inference using *L=*10 (8 given and 2 predicted frames) requires about 2.5 hours; inference using *L=*43 (41 given and 2 predicted frames) requires about 10.5 hours; and inference using *L=*51 (41 given and 10 predicted frames) requires about 12.5 h. However, it should be noted that generating 10 missing frames in steps of 2 (*s=*2) requires 5 consecutive runs, meaning about 5 more time than generating 10 missing frames in a single step.

## V. Conclusion

This paper introduced MW-RaMViD, a diffusion-based method that treats a cryo-ET tilt series as a video frame sequence and performs MW completion by synthesizing the unmeasured high-tilt projections prior to 3D reconstruction. Experiments on synthetic noisy tilt series show that sampling protocol design strongly affects MW completion quality, where generating many frames at once increases error accumulation toward the maximum unknown tilt angle (±90°), while progressive completion with smaller step sizes better preserves structural content. Increasing the number of given frames further improves completion, at the cost of additional runtime. Overall, our results show that MW-RaMViD-based reconstructions consistently outperform the reconstruction obtained from tilt series with an empty MW, and suggest that diffusion probabilistic models provide a promising direction for cryo-ET. The next step is to validate MW-RaMViD on experimental cryo-ET tilt series and assess its impact on downstream tomogram analysis.

## Acknowledgment

We acknowledge the support of the ANR (ANR-23-CE45-0012-03 to SJ), access to HPC resources of CINES and IDRIS granted by GENCI (AD010714089R1 and AD010710998R3 to SJ), and EDITE doctoral school of Sorbonne University (PhD studentship to NH).

